# Synergistic action of actin binding proteins regulate actin network organization and cell shape

**DOI:** 10.1101/2023.09.15.553706

**Authors:** Murielle P. Serres, Matthew B. Smith, Geneviève Lavoie, Philippe P. Roux, Ewa K. Paluch

## Abstract

Animal cell shape is largely determined by the organization of the actin cytoskeleton. Spread shapes result from a balance between protrusive actin networks and contractile stress fibers, while rounded shapes are supported by a contractile actomyosin cortex. The assembly and regulation of distinct types of actin networks have been extensively studied, yet, what determines which networks dominate in a given cell remains unclear. In this Brief Report, we explore the molecular regulation of overall actin organization and resulting cell shape. We use our recently published comparison of the F-actin interactome in spread interphase and rounded mitotic cells to establish a list of candidate regulators of actin networks in spread cells. Utilizing micropatterning and automated image analysis we quantitatively analyze how these candidates affect actin organization. Out of our initial 16 candidates, we identify subsets of proteins promoting stress fibers or regulating their arrangement. Interestingly, no single regulator depletion caused significant cell shape change. However, perturbing two hits simultaneously, supervillin and myosin II, led to stress fiber disassembly and cell rounding. Overall, our systematic investigation shows that actin networks are robust to perturbations, and identifies regulatory modules controlling overall actin organization and resulting cell shape.

## Introduction

Precise control of cell shape is essential for cell physiology, and shape deregulation lies at the heart of many developmental defects and pathological disorders (Heisenberg and Bellaiche, 2013). In animal cells, shape is primarily the result of forces exerted on the plasma membrane by underlying actin networks (Munjal and Lecuit, 2014; Pollard and Cooper, 2009). Actin networks can be arranged in a variety of architectures, which support different cellular shapes (Fletcher and Mullins, 2010). Specifically, spread cell shapes are the result of a balance between protrusive lamellipodia and contractile stress fibers (Burnette et al., 2014; Kumari et al., 2020). Rounded cells, on the other hand, are supported by a an actomyosin cortical network, and contractile tension at the cortex generates intracellular pressure and promotes rounding (Clark et al., 2014; Stewart et al., 2011). Thus, the particular types of network that dominate actin organization largely determine cellular shape (Fletcher and Mullins, 2010).

Cell shape changes, such as transitions between spread and rounded cell morphologies, result from switches in actin organization. Such transitions are displayed in a variety of physiological processes including epithelial-to-mesenchymal transitions (Lamouille et al., 2014), amoeboid-mesenchymal transitions in migrating cells (Paluch et al., 2016), mitotic rounding and post-mitotic spreading (Ramkumar and Baum, 2016), and fate transitions in stem cells (De Belly et al., 2021). To understand how cell shape changes are controlled, it is essential to uncover not only how actin network organization is regulated but also what determines which type of networks dominate actin architecture in a given cell.

The protein composition of specific actin networks, and the role of various actin-binding proteins (ABPs) in the assembly and regulation of these networks have been extensively studied (Chugh and Paluch, 2018; Pollard and Borisy, 2003; Pollard and Cooper, 2009; Tojkander et al., 2012). In contrast, transitions between different types of networks have mostly been investigated from an upstream regulation perspective. The small GTPase Rho has been shown to promote rounded, cortex-dominated cell shapes while Rac has been shown to promote spread, lamellipodia-dominated morphologies (Sanz-Moreno and Marshall, 2010). However, most cellular actin networks comprise over 100 distinct ABPs (Liu et al., 2022; Obermann et al., 2019; Vadnjal et al., 2022) and how exactly small GTPases control downstream actin network composition and organization remains poorly understood. It has recently been proposed that transitions between lamellipodia and filopodia are governed by competition for actin monomers by different actin nucleators, with the Arp2/3 complex promoting lamellipodia and formins favoring filopodia (reviewed in (Suarez and Kovar, 2016)). However, both the Arp2/3 complex and formins contribute to cortical actin nucleation (Bovellan et al., 2014). Thus, interfering with the activity of specific actin nucleators alone does not lead to transitions between lamellipodial/filopodial and cortical networks that drive cell spreading or rounding (Bovellan et al., 2014; Cao et al., 2020; Chugh et al., 2017). Overall, which of the multitude of cellular ABPs specify the dominating actin networks, and thus the resulting cell shape, at a given time remains unclear.

In a recent study, we directly compared the “F-actin interactome” (the ABPs binding actin filaments) in spread interphase cells, where actin is mostly organized into lamellipodia and stress fibers, and rounded mitotic cells, where cortical actin dominates (Serres et al., 2020). The ABPs found to be enriched in the F-actin-binding fraction from interphase cells constitute potential candidates for the regulation of actin organization promoting spread shapes. In this Brief Report, we used this list of ABPs enriched in the spread cells F-actin interactome to investigate how interphase actin organization, and resulting cell shapes, are regulated (**Figure 1A**). We first developed a pipeline for quantitative analysis of actin organization, and identified proteins controlling stress fiber formation and distribution. This allowed us to identify subgroups of proteins that regulate stress fiber organization, or promote stress fibers at the expense of other networks. While no depletion of any single candidate triggered the rounding of spread cells, we showed that simultaneously perturbing two of the key hits, supervillin and myosin II, leads to stress fiber disassembly and cell rounding. Our findings show that modulating the activity of small subsets amongst the myriad of cellular ABPs can induce transitions between actin architectures and, as a result, drive cell shape transitions.

**Figure 1:**
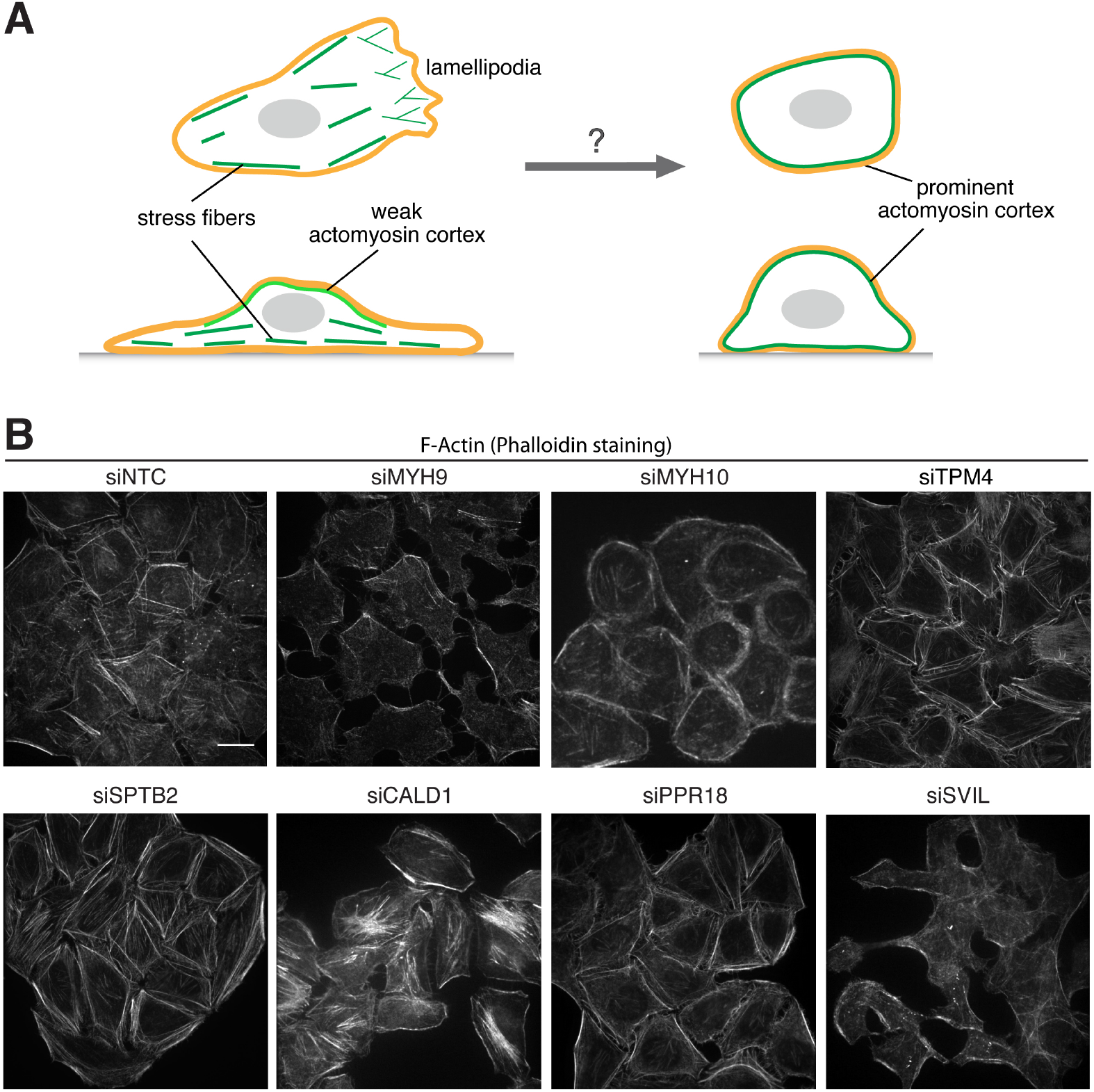
Regulation of stress fiber actin organization. **(A)** Framework of the study: Spread cell shape is dominated by lamellipodia and stress fibers while rounded morphologies are supported by a strong actomyosin cortex. We investigate which ABPs regulate overall actin organization in spread cells, and as a result, cell shape. **(B)** Representative images of cells depleted with siRNA control (siNTC) or with different siRNA targeting candidate regulators. Cells were labelled with phalloidin to stain F-actin. Scale bar, 20 µm. See Figure S1B for the full set of experimental conditions.

## Results and discussion

### Investigating candidate regulators of interphase actin organization

We set out to investigate which ABPs promote the formation of the actin networks supporting spread cell shapes at the expense of the actomyosin cortex that supports rounded shapes (**Figure 1A**). As candidate proteins, we used the list of ABPs enriched in interphase spread cells compared to rounded mitotic cells identified in our recent systematic investigation of the F-actin interactome (Serres et al., 2020). Out of the 26 candidates identified in the F-actin-interactome analysis (Table 1 in (Serres et al., 2020)), we narrowed down the candidate list and focused on 16 proteins that have been shown to directly interact with actin (**Table 1**). We then asked how depletion of these candidates affects actin network organization in spread interphase HeLa cells. Target proteins were depleted using siRNA, and mRNA transcript depletion efficiency was confirmed using qPCR (**Figure S1A**). F-actin distribution was then visually assessed using phalloidin staining. We observed some differences in F-actin distribution, with some treatments resulting in a clearly more prominent stress fiber network (e.g. CALD1 siRNA), and others reducing stress fibers (e.g. MYH9 siRNA) (**Figures 1B and S1B**). However, interphase cells display a variety of shapes and as a result, F-actin distribution and intensity appeared heterogenous even under control conditions (siNTC panels in **Figures 1B and S1B**), making a quantitative comparison of actin networks between conditions challenging. Thus, though depletion of some candidate regulators seemed to affect interphase actin networks, cell shape heterogeneity precluded objective quantifications and conclusions.

**Table 1:**
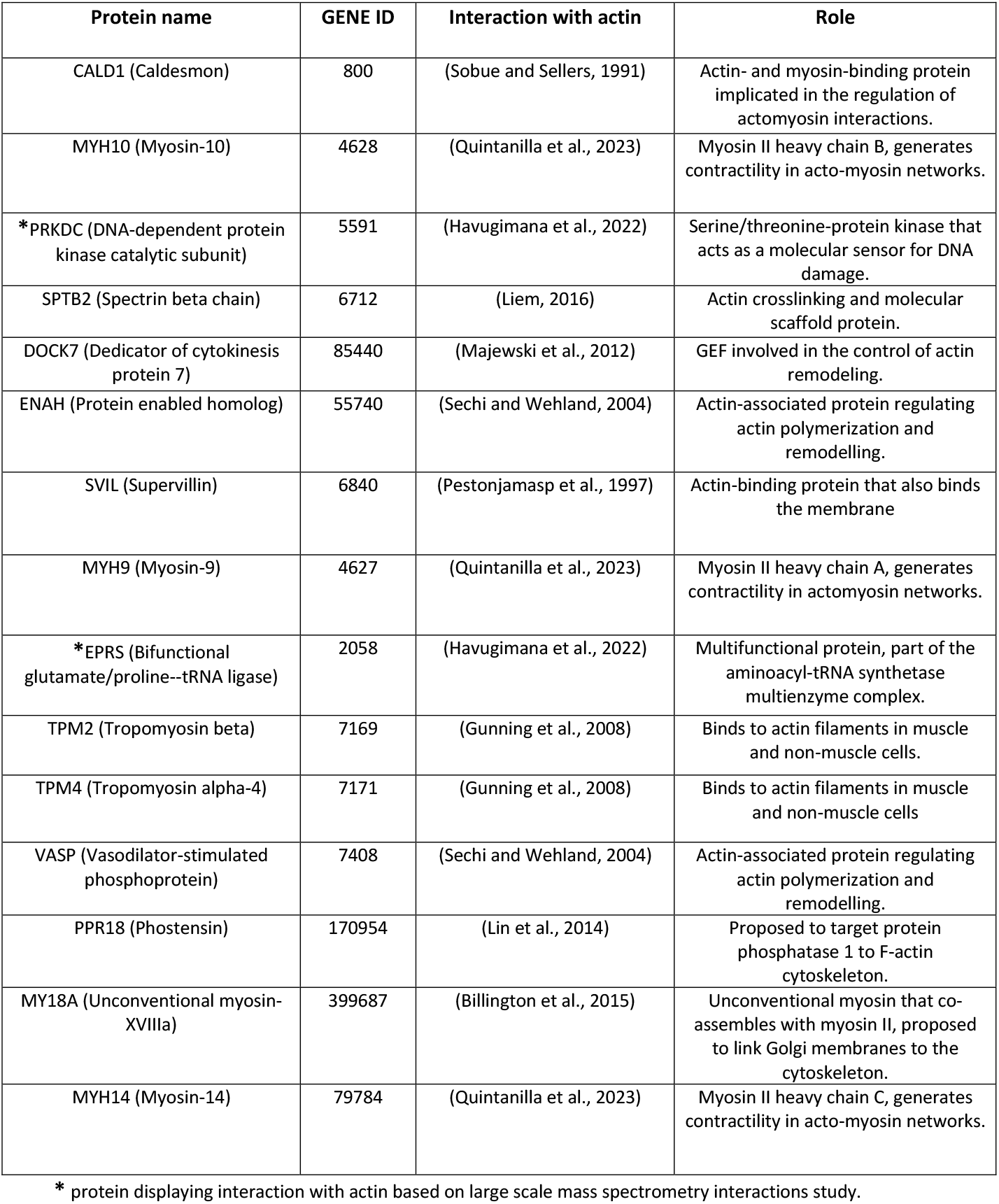
List of candidates identified in the analysis of the F-actin interactome as enriched in spread interphase cells compared to rounded mitotic cells, and known to directly interact with actin.

### Quantitative analysis of actin organization upon candidate depletion using micropatterns to standardize actin distribution

To avoid confounding effects resulting from the inherent variability in cell shape both within and between experimental conditions, we plated cells on fibronectin-coated micropatterns, which standardize cell shape and F-actin distribution across cell populations, facilitating quantitative comparisons (Théry et al., 2006) (**Figures 2, S2 and S3**). We used H-shaped patterns, where interphase cells display prominent peripheral stress fibers on the non-adhesive edges of the patterns, as previously described ((Théry et al., 2006); **Figure 2A**). After depletion of each candidate protein, cells were detached and plated on fibronectin patterns for 3 hours before fixation. Cells were then stained with phalloidin to label F-actin, and phalloidin distribution was quantified using custom-written software (**Figure 2A-C**). To visualize changes in actin distribution, we also generated average actin distributions, following an approach developed in (Théry et al., 2006) (**Figures 2, D and E, S2A and B**). We observed that none of the protein depletions tested led to a complete loss of stress fibers, suggesting that overall interphase actin architecture is robust to perturbations by one single factor (**Figure 2, D and E**). To quantitatively analyse actin distribution upon candidate depletion, we measured two parameters: R_1_, which compares the intensity of phalloidin staining at the periphery versus the central region of a cell, and R2, which compares phalloidin intensity at the non-adhesive versus adhesive edges of a cell (**Figure 2, A and B**). A decrease in R_1_ corresponds to a reduction of actin stress fibers at the cell periphery, whereas changes in R_2_ correspond to changes in the organization and distribution of peripheral stress fibers. We considered R_1_ as the primary readout of the extent to which stress fibers are the dominant actin network in a given condition.

**Figure 2:**
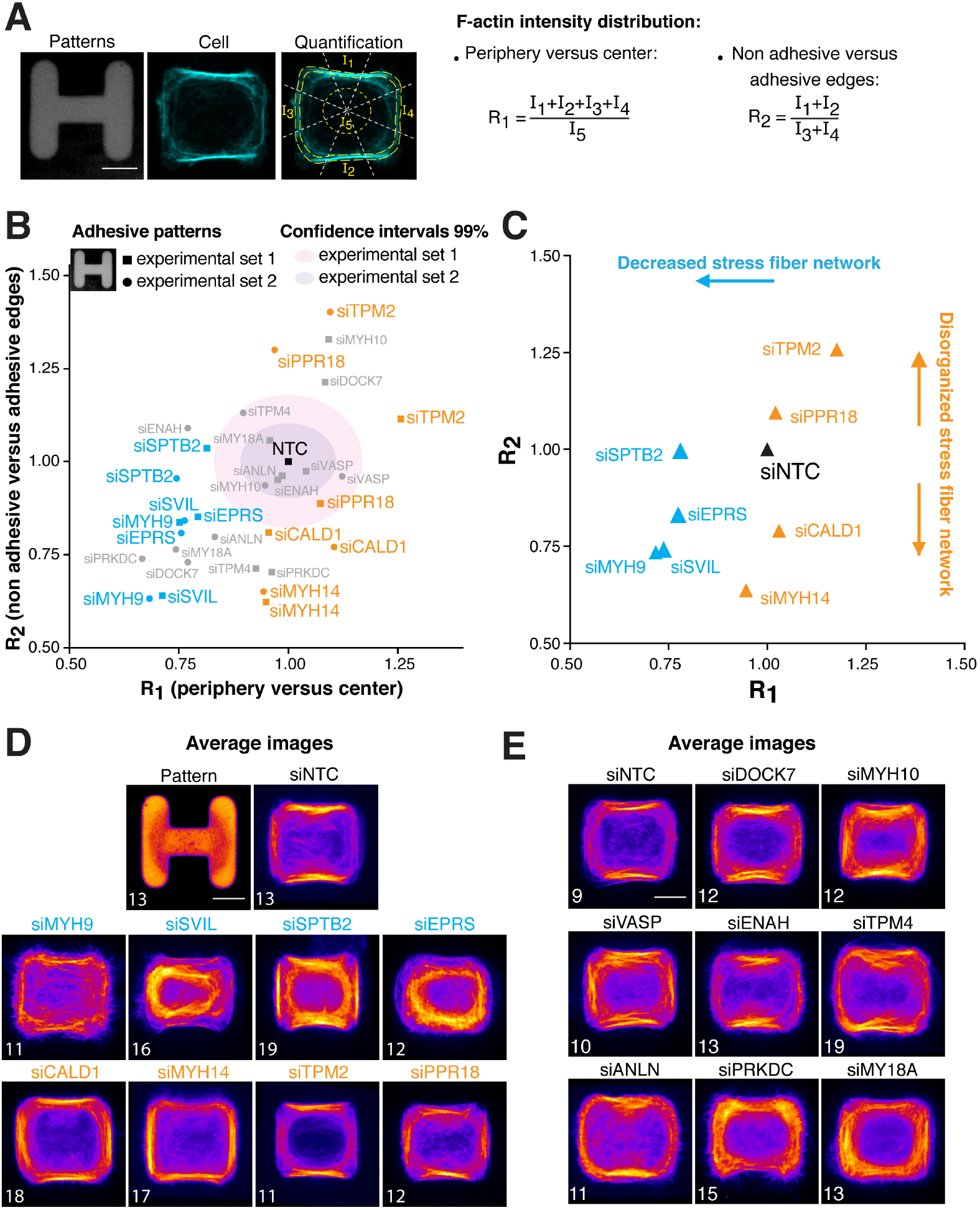
Regulation of stress fiber actin organization in cells on fibronectin micropatterns. **(A)** Representative image of a control cell (middle, phalloidin-labelled F-actin) spread on an H-shaped fibronectin-coated micropattern (left image). Prominent stress fibers form on the non-adhesive edges of the pattern. The right panel illustrates the quantification of actin distribution in cells on micropatterns. The phalloidin staining is used for cell segmentation, mean intensity is then measured in areas at the cell periphery (I_1_ and I_2_ on non-adhesive pattern edges and I_3_ and I_4_ on adhesive edges) and in the cell center (I_5_). The parameters R_1_ and R_2_ characterize actin intensity distributions. Scale bar, 10 µm. **(B, C)** Quantification of the effects of siRNA depletions on F-actin intensity distribution in cells on H-shaped micropatterns, as quantified by the coefficients R_1_ and R_2_ described in **(A)**. R_1_ lower than control corresponds to decreased stress fibers; high R_1_ corresponds to enhanced stress fibers; changes in R_2_ reflect changes in the organization and distribution of the stress fibers network. R_1_ and R_2_ have been normalized to the corresponding control conditions, siNTC, for each experimental set (2 independent experiments). **(B)** Quantification of R_1_ and R_2_ for all the experimental conditions. The pink shaded regions correspond to the 99% confidence intervals of the NTC controls of the two independent experiments. Grey: conditions that fall within the confidence interval in at least 1 experiment, or for which the 2 experiments gave inconsistent results. Blue: conditions resulting in decreased stress fibers; orange: conditions resulting in disorganization of the stress fiber network. **(C)** Mean R_1_ and R_2_ for the conditions significantly perturbing actin distribution. Blue: conditions resulting in decreased stress fibers; orange: conditions resulting in disorganization of the stress fiber network. Each dot represents the mean of the two independent experiments (siNTC (Non-targeting control) n=55, siSVIL n=28, siEPRS n=26, siMYH9 n=16, siSPTB2 n=35, siTPM2 n=25, siCALD1 n=37, siMYH14 n=27, siPPR18 n=23). **(D, E)** Average F-actin intensity distributions for cells on H-shaped micropatterns for the conditions displayed in panel **B. (D)** Images for the conditions perturbing actin distribution displayed in panel **C. (E)** Images corresponding to the grey data points in panel **B**. The number of cells averaged is displayed on the left bottom of each picture. Scale bars, 10 µm.

We quantified actin distribution in cells depleted for the 16 candidate proteins in two independent experiments, and analyzed how candidate depletion affected the R_1_ and R_2_ parameters (**Figure 2B**, R_1_ and R_2_ were normalized to the corresponding NTC values). Conditions which were not statistically different from controls, or for which the changes in R_1_ and R_2_ were inconsistent between the two experiments, were excluded from further analysis. This analysis identified 8 candidate proteins for which depletion significantly and consistently affected actin distribution in interphase cells plated on adhesive micropatterns (**Figure 2C**). These 8 hits could be subdivided into two groups (**Figure 2, B and C**). The depletion of myosin heavy chain IIA (MYH9), supervillin (SVIL), spectrin (SPTB2), and the tRNA ligase EPRS all led to a reduction in stress fibers (**Figures 2B-D** and **S2A**, blue data points). This reduction was associated with the formation of a less organized, cortex-like actin network at the cell membrane in the central region of the cell (**Figure 2D**). In contrast, the depletion of caldesmon (CALD1), myosin heavy chain IIC (MYH14), tropomyosin 2 (TPM2) and the F-actin cytoskeleton-targeting subunit of protein phosphatase 1 (PPR18) affected stress fiber distribution (**Figures 2B-D** and **S2A**, orange data points). In particular, depletion of MYH14 and CALD1 led to a redistribution of stress fibers from the non-adhesive cell edges to the entire cell periphery.

Analysis of actin distribution on a differently shaped H-pattern, with a larger adhesive region, identified the same two subgroups of regulators of stress fiber organization, supporting the validity of our findings (**Figure S3A-D**). This additional experimental condition identified further regulators, with depletion of the kinase protein (PRKDC) leading to a reduction in stress fibers, and depletion of tropomyosin 4 (TPM4) and Anillin (ANLN) leading to changes in stress fiber distribution.

Together, our analysis identifies a key set of ABPs that display an increased association with F-actin in spread interphase cells (Serres et al., 2020) and that regulate actin distribution and organization in these cells. Some of the regulators identified display functions consistent with previous reports. For instance, the effect of MYH9 is consistent previous observations that MYH9 depletion, or the inhibition of myosin II activity by blebbistatin, reduce stress fibers in a variety of spread cells (Cai et al., 2006; Hotulainen and Lappalainen, 2006). Caldesmon and various tropomyosins have previously been involved in regulating stress fiber network organization (Helfman et al., 1999; Kordowska et al., 2006; Tojkander et al., 2011). For instance, tropomyosin depletion has been shown to change the organization of actin stress fibers, leading to more organized but less contractile fibers in rat embryonic fibroblasts (Hu et al., 2019). Caldesmon knockout in U2OS cells plated on micropatterns led to a less regularly organized F-actin network and a redistribution of stress fibers away from non-adhesive pattern edges (Kokate et al., 2022), in line with our findings. Taken together, this consistency with previous reports supports the validity of our approach. Our quantitative analysis allowed for a systematic investigation of the regulation of interphase actin organization, identifying subgroups of ABPs controling either stress fiber organization or formation.

### Supervillin depletion combined with myosin II inhibition lead to actin reorganization and cell rounding

Next, we investigated whether depletion of the identified regulators of interphase actin organization would also elicit a transition between spread and rounded cell shapes. We focused on the 4 ABPs for which depletion led to a reduction in stress fibers (**blue data points on Figure 2B-D**) and asked whether depleting them could facilitate a transition to a rounder cell morphology. We excluded EPRS from our analysis since, even though it has a putative actin-binding domain (Carninci et al., 2005), it is not known to directly regulate actin. As cell shape is constrained on micropatterns, we depleted candidate regulators in cells plated in regular glass-bottom dishes, where shape is not constrained. We found that neither myosin heavy chain IIA (MYH9) nor spectrin B2 (SPTB2) depletion had any significant impact on cell shape, and supervillin depletion appeared to lead to slightly more elongated cell morphologies (**Figure 3A**, T0 left-most images and **Figure 1**). We then asked whether significant actin reorganization needed for rounding required the action of more than one factor. We thus inhibited the activity of myosin II using blebbistatin in supervillin- and spectrin-depleted cells. We checked that blebbistatin treatment led to a reduction of stress fibers similar to the effect of MYH9 depletion in cells plated on micropatterns (**Figures S2C and S3E**). Blebbistatin addition had no significant effect on cell shape in control and spectrin-depleted cells (**Figure 3A-C**). However, we observed that addition of blebbistatin in supervillin-depleted cells led to significant cell rounding and a transition from stress-fiber-dominated to a more cortical actin organization (**Figures 3D and S4A)**. To verify that the cell rounding observed was not due to, by chance, imaging cells entering mitosis, we monitored DNA condensation as a readout of mitotic entry. We could observe cell rounding upon blebbistatin addition to SVIL-depleted cells without any sign of DNA condensation, confirming the cells displayed rounding in interphase (**Figure S4B**). Interestingly, the extent of interphase rounding was comparable to the beginning of mitotic cell rounding (**Figure S4C**). Taken together, these data indicate that depletion of the membrane-associated protein supervillin combined with inhibition of myosin II activity is sufficient to induce actin reorganization and a transition from spread to rounded cell morphology.

**Figure 3:**
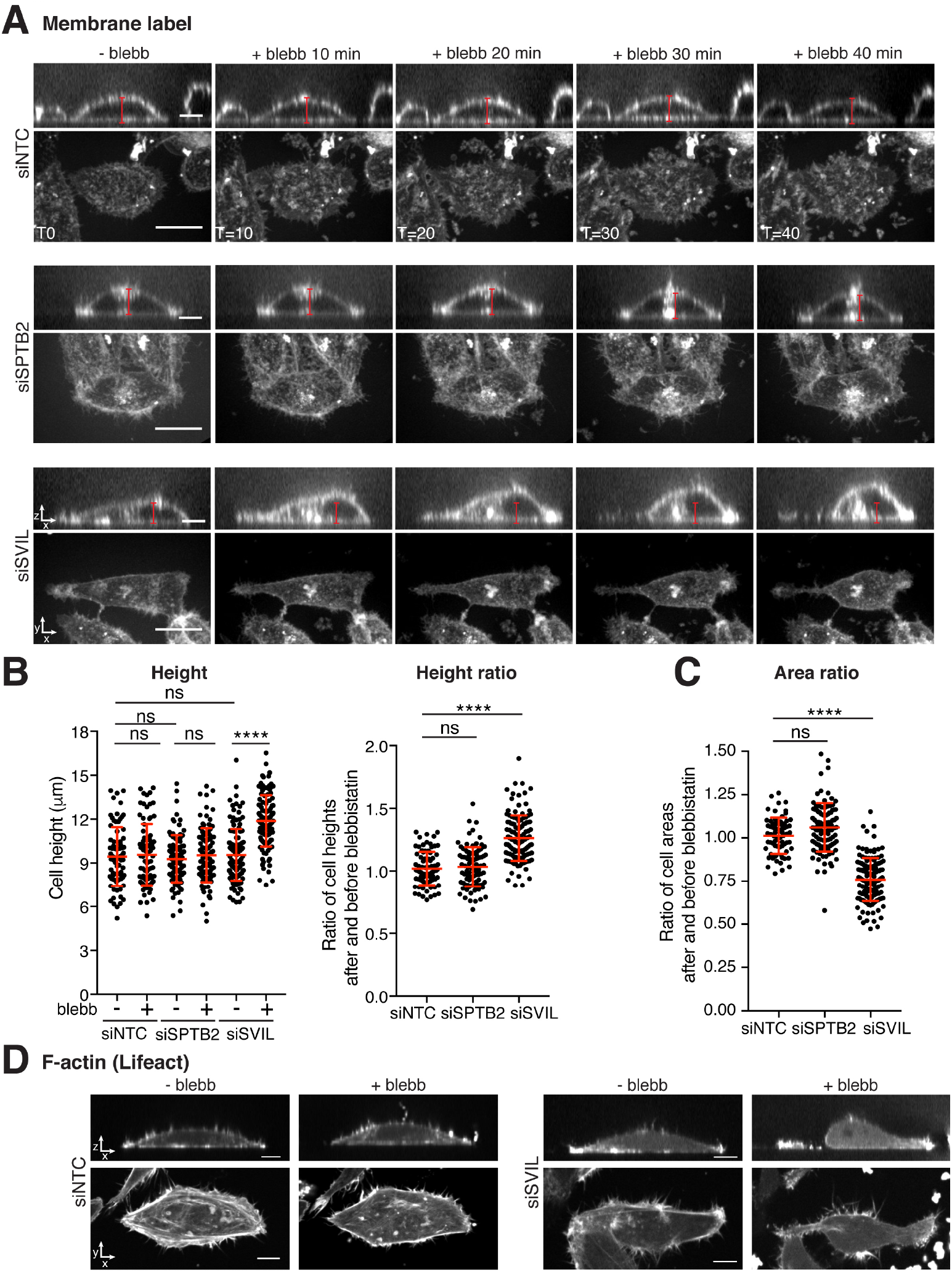
Combining supervillin depletion with myosin II inhibition leads to cell rounding in interphase. **(A)** Representative time lapse images of cells treated with non-targeting control siRNA (siNTC, top), siRNA against spectrin (siSPTB2), siRNA against supervillin (siSVIL), and treated with the myosin II inhibitor blebbistatin (blebb.) at t=0. Far red cell mask was used to label the membrane. Scale bars XY, 20 µm, XZ 10 µm. **(B)** Cell height and cell height ratio in control and supervillin-depleted cells before and 40 min after the addition of blebbistatin. ANOVA non-parametric Kruskal-Wallis test; P-values: **** P<0.0001, ns: non-significant. **(C)** Ratio of projected cell areas 40 min after blebbistatin addition and before treatment, in control cells and supervillin-depleted cells. ANOVA non-parametric Kruskal-Wallis test; P-values: **** P<0.0001, ns: non-significant (P=0.41). **(D)** Representative images of control and supervillin-depleted cells expressing GFP-Lifeact before and after addition of blebbistatin. Scale bars, 10 µm.

Taken together, our quantitative analysis of actin organization, using adhesive micro-patterns to standardize cell morphology and actin distribution identified a subgroup of ABPs, for which depletion affects peripheral stress fiber organization, and another subgroup of ABPs, for which siRNA depletion reduces cellular stress fibers (**Figure 2**). Interestingly, none of the individual treatments led to a complete transition from stress fibers to a cortical network. This strongly suggests that interphase actin organization is the result of a complex interplay of multiple ABPs acting together with redundancy in function. However, we could show that depletion of the membrane-associated scaffolding protein supervillin in combination with inhibition of myosin II activity leads to actin reorganization towards a more cortex-like network, and cellular rounding (**Figure 3**). Supervillin, which binds both actin and myosin II, has previously been shown to enhance myosin activity and promote peripheral stress fiber-like actin bundles (Takizawa et al., 2007). Together with our data, this indicates that even though myosin II is essential for contractility generation in both stress fibers and in the actomyosin cortex, it also plays a central role in promoting stress fibers at the expense of the cortex.

In summary, our investigation of actin organization as a whole, rather than focusing on a specific type of network, unveils key regulators of overall actin organization and cell morphology. As such, our study paves the way to a molecular understanding of shape changes in other biological contexts where cells transition between spread and rounded morphologies.

## Acknowldgements

We thank members of the Paluch lab, especially R. Peters, J. Lefévère-Laoide and D. Bodor, for feedback on the project and manuscript, and K. Chalut for feedback on the manuscript. We thank W. Pönisch and A. Stubb for help with data analysis. We thank A. Vaughan and the LMCB Light Microscopy Facility for support with microscopy, and C. Dix (B. Baum lab) for technical help with generating adhesive micropatterns. We acknowledge support from the Medical Research Council UK (MRC programme award MC_UU_12018/5 to E.K.P.), the Human Frontier Science Program (Young Investigator Grant RGY 66/2013 to P.P.R. and E.K.P.), the European Research Council (Starting Grant 311637-MorphoCorDiv and Consolidator Grant 820188-NanoMechShape to E.K.P.), an operating grant from the Canadian Institute of Health Research (MOP-142374 and PJT-178154) and a Cancer Research Society grant (840382) to P.P.R..

## Authors contributions

M.P.S and E.K.P designed the research; M.P.S carried out most of the experiments and analyzed the data; M.B.S developed analysis tools for micropattern analysis; G.L. carried out the qPCR experiment; M.P.S and E.K.P wrote the paper; P.P.R. provided technical guidance. Funding Acquisition: P.P.R, E.K.P. All authors discussed the results and manuscript.

## Methods

### Cell lines, cell culture, drug treatments and transfections

All cell lines were grown at 37°C with 5% CO_2_ in DMEM GlutaMAX, 4.5 g/L glucose (Gibco, Invitrogen/ Life Technologies) supplemented with 10% fetal bovine serum (Sigma), 1% penicillin-streptomycin (Gibco, Invitrogen/ Life Technologies). HeLa Kyoto cells were obtained from the Research Institute of Molecular Pathology (Vienna, Austria). HeLa Kyoto cells stably expressing H2B-mCherry and GFP-Lifeact were described in (Serres et al., 2020). For live imaging, cells were labelled with CellMaskTM Deep Red plasma membrane Stain (Thermo Fisher Scientific). For myosin II inhibition, blebbistatin was used at 100 μM. For protein depletion, RNAi max lipofectamine (Invitrogen) reverse transfection was used for transfection according to the manufacturer’s instructions. siRNA SMARTpool from Dharmacon were used at a final concentration of 30 nM for all the experiments.

### Immunofluorescence

For fibronectin micropattern experiments, siRNA-depleted cells were synchronized in interphase with thymidine (Sigma) at 2 mM for 22 h and were washed 2 times with PBS then detached using versene and seeded onto micropatterned coverslips. After 3 hours, cells were permeabilized for 15 s with 0.5% Triton X-100 in Cytoskeleton buffer pH 7.4 (10 mM MES, 138 mM KCl, 3 mM MgCl_2_, 2 mM EGTA) supplemented with 4.51% of sucrose to adjust osmotic pressure (Cytoskeleton buffer with sucrose; CBS) and then fixed in 3% PFA in CBS for 15 min. Cells were then washed 3 times in CB and incubated for 45 min in CB with phalloidin conjugated with Alexa 488 (Life Technology A12379). After 3 washes in CB, coverslips were mounted using Vectashield (Vectorlabs).

### Adhesive micropattern preparation

Square coverslips (No 1.5, 170 nm) were washed with 1 M HCl for 15 min and air dried. Coverslips were then activated with plasma treatment and incubated with 0.1 mg/mL of fluorescent PLL-g-PEG solution (PLL(20)-g[3.5]-PEG(2)/Atto663; SurfaceSolutions GmbH, Zurich) in 10 mM Hepes pH 7.4 for 30 min at room temperature. Coverslips were placed onto a quartz photomask designed with specific shape patterns (Delta Mask, The Netherlands) and illuminated with UV light for 5 min. The illuminated surface was incubated with 25 µg/mL of fibronectin in 100 mM NaHCO_3_ (pH 8.5). Micropatterned coverslips were washed with PBS before use.

### Quantification of micropattern experiments

Images were taken using a Nikon spinning disk microscope; 2 μm Z stack with ΔZ = 0.2 μm were acquired at the bottom of the cell and analyzed using custom-written ImageJ plugins. First, images were cropped and rotated based on the H-shaped micropatterns, so that all the micropatterns overlapped (see micropattern average images in **Figures 2D, S2C, S3C and E**). Cells were then segmented using an active contour-based algorithm. Briefly, segmentation involved a maximum projection of the phalloidin channel, a k-means threshold and connected components algorithms to separate the cell from the background, and cell border extraction as an active contour. The active contours were then deformed using JFilament (Smith et al., 2010) to attract to the maximum gradient, refining the segmentation, and manually improved where the automatic locally failed. For measurements of intensity at the center of the cell, a circle containing one third of the cell area and centered at the cell center of mass was generated (region I_5_ in **Figure 2A**), and mean intensity within this circle was measured. For measurements of F-actin intensity at the cell periphery, the cell contour was divided into 8 sections of equal angular size from the cell center, and the mean intensity in a 15 pixel wide region along the contour was measured in the section corresponding to non-adhesive and adhesive cell edges (regions I_1_ to I_4_ in **Figure 2A**). Only pixel values that exceeded the inner mean intensity, I_5_, were included in the average. The analysis ImageJ plugin will be made freely available online.

### RNA extraction and RT-qPCR

Total RNA from Hela Kyoto cells was extracted using RNeasy Mini Kit (Qiagen) 48 h after siRNA transfection. Reverse transcription and qPCR were performed as previously described (Chugh et al., 2017). List of the primers used:

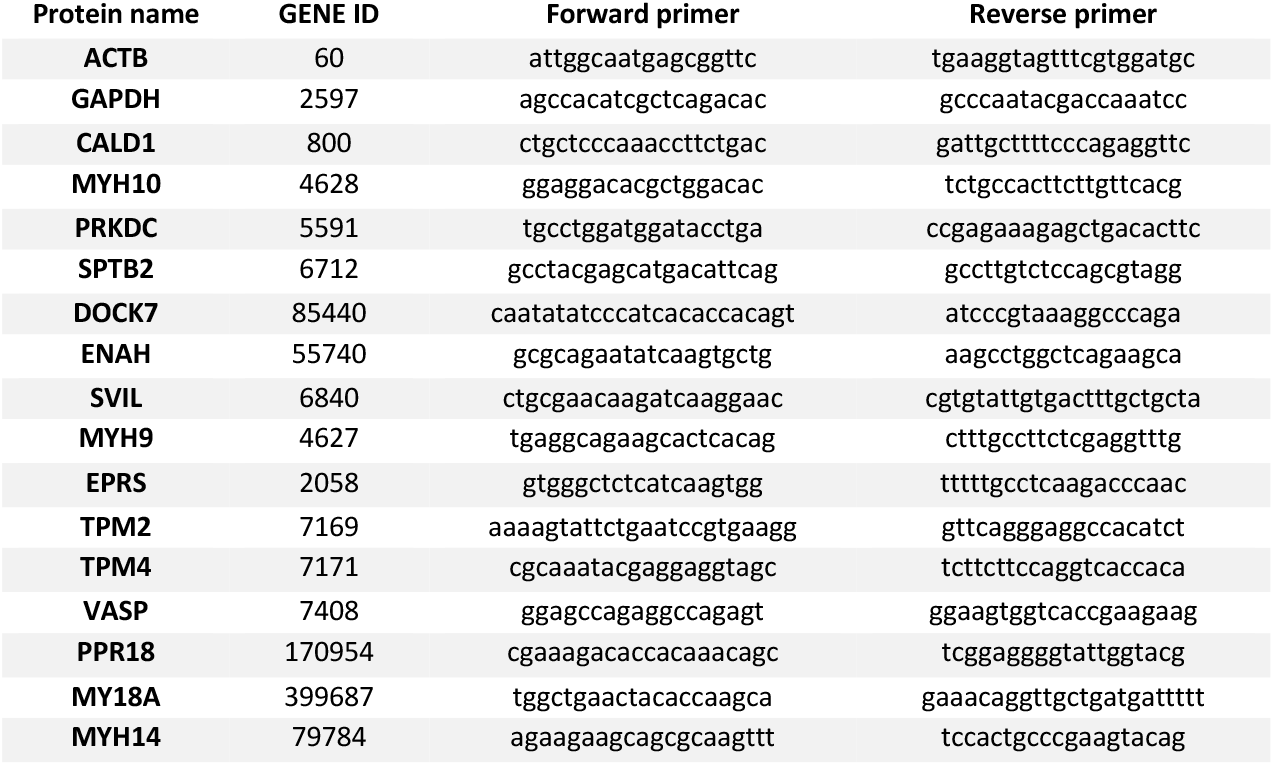

### Live cell imaging

Cells were imaged in phenol-red free and CO_2_-independent medium (Invitrogen) at 37°C using a Nikon TE2000 microscope equipped with a Plan Fluor x60/1.4 DIC H objective (Nikon), a PerkinElmer ERS Spinning disk system, a Digital CCD C4742-80-12AG camera (Hamamatsu), and controlled by Volocity 6.0.1 software. Stacks of 25 z-planes 1 μm apart were acquired every 10 min. Cell areas were measured on projected z-stacks and cell heights were extracted from intensity profiles of a plasma membrane label along the z direction (peak-to-peak distance in the intensity profile) using Fiji software.

### Statistical Analysis

Graphs and statistical tests were produced in GraphPad Prism. Details of the statistical tests used, exact value of n, definition of error bars on graphs, and number of experiments performed are detailed in the figure legends.

