## Supplemental figures for "Synergistic action of actin binding proteins regulate actin network organization and cell shape"

4 Supplemental Figures

### Figure S1

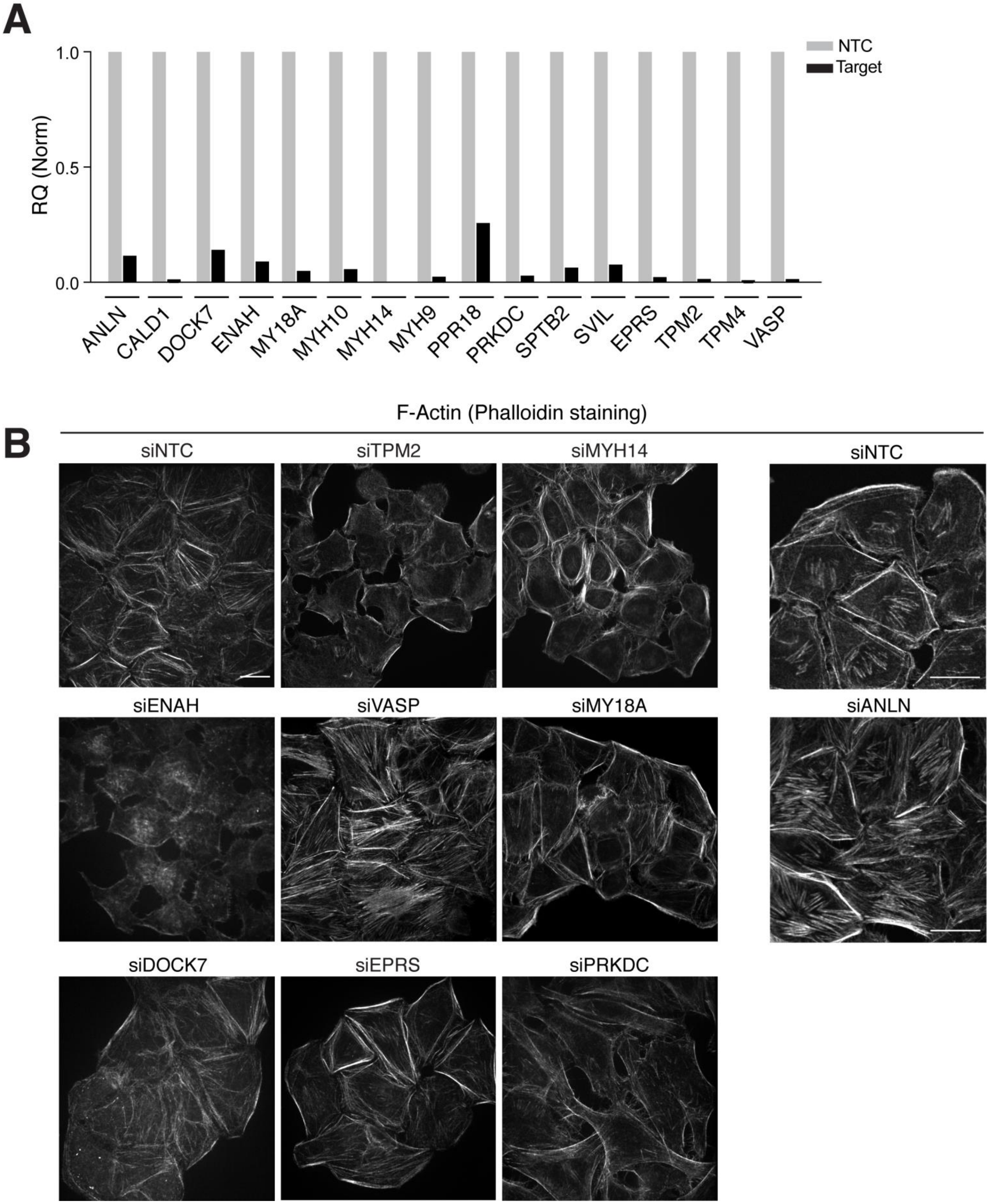
A
NTC
Target
RQ (Norm)
1.0
0.5
0.0
ANLN
CALD1
DOCK7
ENAH
MY18A
MYH10
MYH14
MYH9
PPR18
PRKDC
SPTB2
SVIL
EPRS
TPM2
TPM4
VASP
B
F-Actin (Phalloidin staining)
siNTC
siTPM2
siMYH14
siNTC
siENAH
siVASP
siMY18A
siANLN
siDOCK7
siEPRS
siPRKDC

**Figure S1: Additional experiments on the effects of siRNA depletion on F-actin organization, related to Figure 1.**

**(A)** mRNA expression levels (RQs) in cells transfected with siRNAs against the different interphase hits. Target RQs were normalized to the RQs of the corresponding non-targeting control (NTC) siRNAs. **(B)** Representative images of cells depleted with siRNA control (siNTC) or with different siRNA targeting candidate regulators. Cells were labeled with phalloidin to stain F-actin. Note that the imaging of siANLN-treated cells and associated control was performed with different parrameters and, as a result, the images displayed have a slightly different magnification compared to other conditions. Scale bars, 20 µm.

### Figure S2

**A** **Average images**

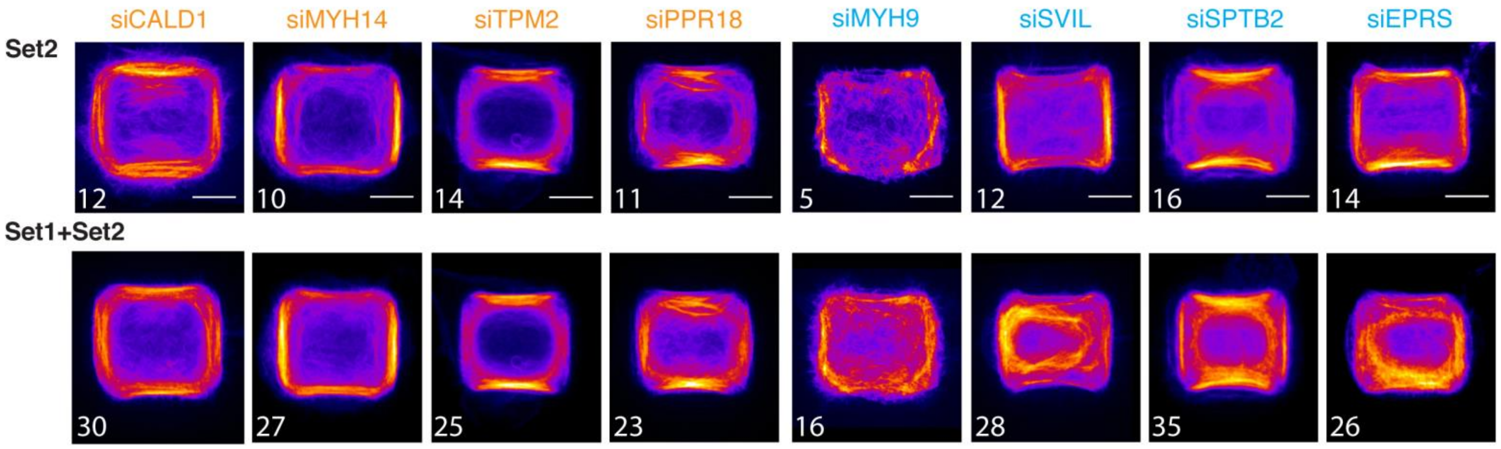


**B** **Average images**

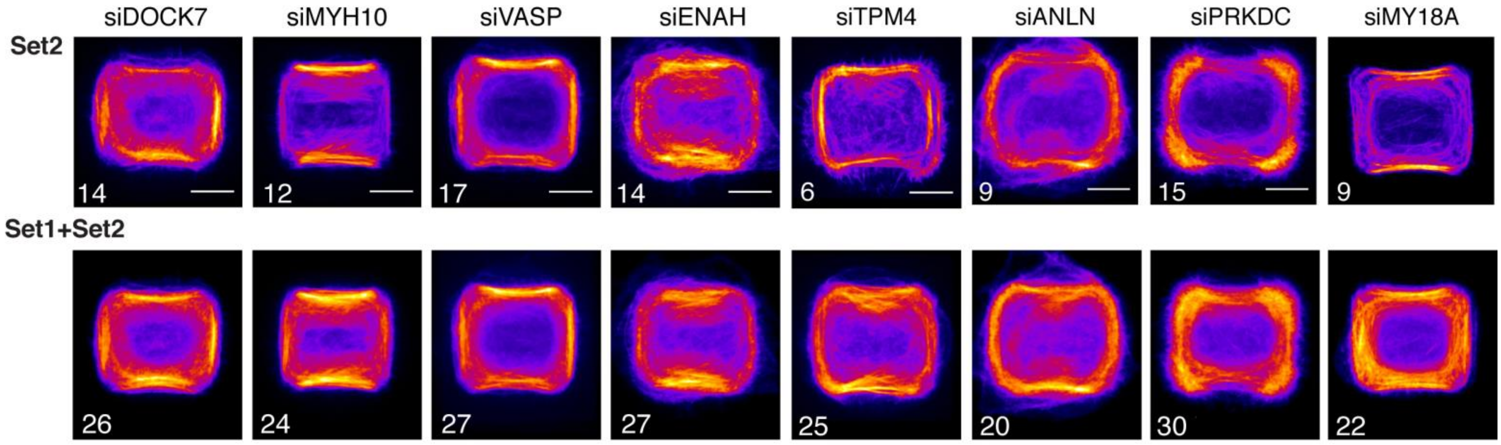


**C** **Average images**

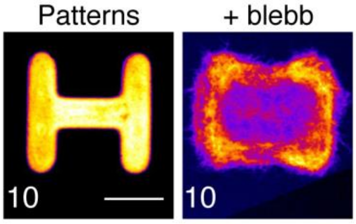

**Figure S2: Additional experiments on the regulation of actin organization in cells on thin H-shaped micropatterns, related to Figure 2.**

**(A, B)** Average F-actin intensity distributions for cells on H-shaped micropatterns. Images corresponding to the experimental Set 2 (Set 1 is displayed in Figure 2D and E), and to the average of the 2 experimental sets (Set1+Set2). **(A)** Images for the conditions perturbing actin organization identified in Figure 2. **(B)** Images for the conditions that do not significantly perturb actin organization, corresponding to the grey data points in Figure 2B. The number of cells averaged is displayed on the left bottom corner of each image. Scale bars, 10 µm. **(C)** Average F-actin intensity distribution for cells treated with blebbistatin on thin H-shaped micropatterns. The number of cells averaged is indicated on the image. Scale bar, 10 µm.

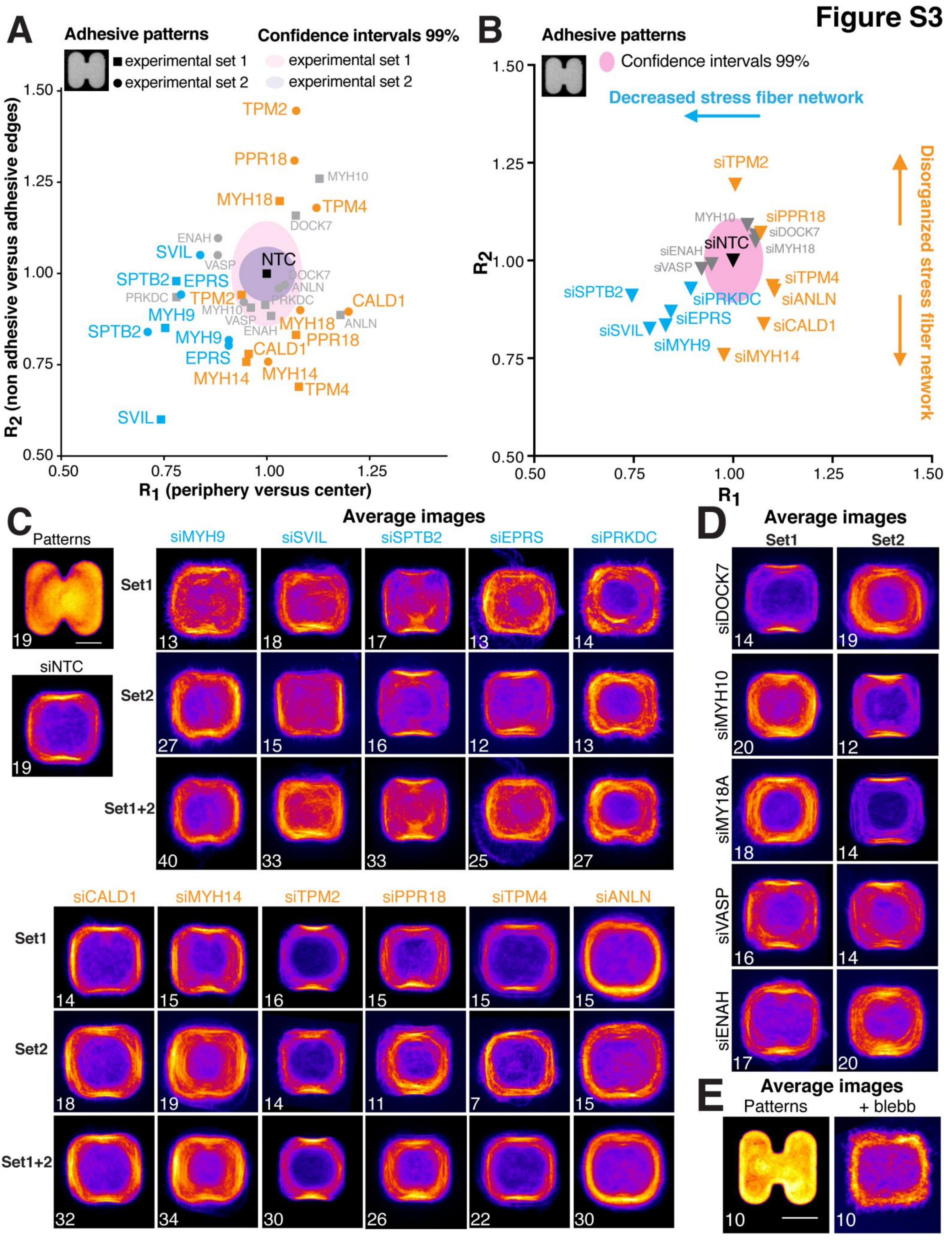

Figure S3
A
Adhesive patterns
experimental set 1
experimental set 2
Confidence intervals 99%
experimental set 1
experimental set 2
R2 (non adhesive versus adhesive edges)
R1 (periphery versus center)
TPM2
PPR18
MYH10
MYH18
TPM4
DOCK7
ENAH
SVIL
VASP
NTC
SPTB2
EPRS
PRKDC
ANLN
CALD1
MYH9
MYH14
B
Decreased stress fiber network
Disorganized stress fiber network
siTPM2
siPPR18
siDOCK7
siMYH18
siNTC
siENAH
siVASP
siTPM4
siANLN
siSPTB2
siPRKDC
siEPRS
siCALD1
siSVIL
siMYH9
siMYH14
R1
R2
C
Patterns
Average images
siMYH9
siSVIL
siSPTB2
siEPRS
siPRKDC
siNTC
Set1
Set2
Set1+2
siCALD1
siMYH14
siTPM2
siPPR18
siTPM4
siANLN
D
Average images
Set1
Set2
siDOCK7
siMYH10
siMY18A
siVASP
siENAH
E
Average images
Patterns
+ blebb

**Figure S3: Additional experiments on the regulation of stress fiber actin organization in cells on thick H-shaped micropatterns, related to Figure 2.**

**(A, B)** Quantification of the effects of siRNA depletions on F-actin intensity distribution in cells on thick H-shaped micropatterns (pictured top left), as quantified by the coefficients $R_1$ and $R_2$ described in **Figure 2A**. $R_1$ and $R_2$ have been normalized to the corresponding control conditions, siNTC, for each experimental set (2 independent experiments). The pink shaded regions corresponds to the 99% confidence interval of the NTC controls. Grey: conditions that fall within the confidence interval. Blue: conditions resulting in decreased stress fibers; orange: conditions resulting in disorganization of the stress fiber network. Panel **(A)** displays mean $R_1$ and $R_2$ for each experimental set separately, panel **(B)** displays overall mean values for the two sets combined. (siNTC n=94, siSVIL n=33, siEPRS n=25, siMYH9 n=40, siTPM2 n=30, siCALD1 n=44, siMYH14 n=34, siSPTB2 n=33, siPPR18 n=26, siTPM4 n=23, siANLN n=30, siPRKDC n=27, siENAH n=37, siVASP n=30, siMYH18 n=33, siMYH10 n=33, siDOCK7 n=33). **(C, D)** Average F-actin intensity distributions for cells on thick H-shaped micropatterns. Images corresponding to both experimental sets and to the average of the 2 sets (Set1+Set2) are displayed. **(C)** Images for the conditions perturbing actin organization. **(D)** Images for the conditions that do not significantly perturb actin organization, corresponding to the grey data points in Figure S2B. The number of cells averaged is displayed on the left bottom corner of each image. Scale bar, 10 µm. **(E)** Average F-actin intensity distribution for cells treated with blebbistatin on thick H-shaped micropatterns. The number of cells averaged is indicated on the image. Scale bar, 10 µm.

### Figure S4

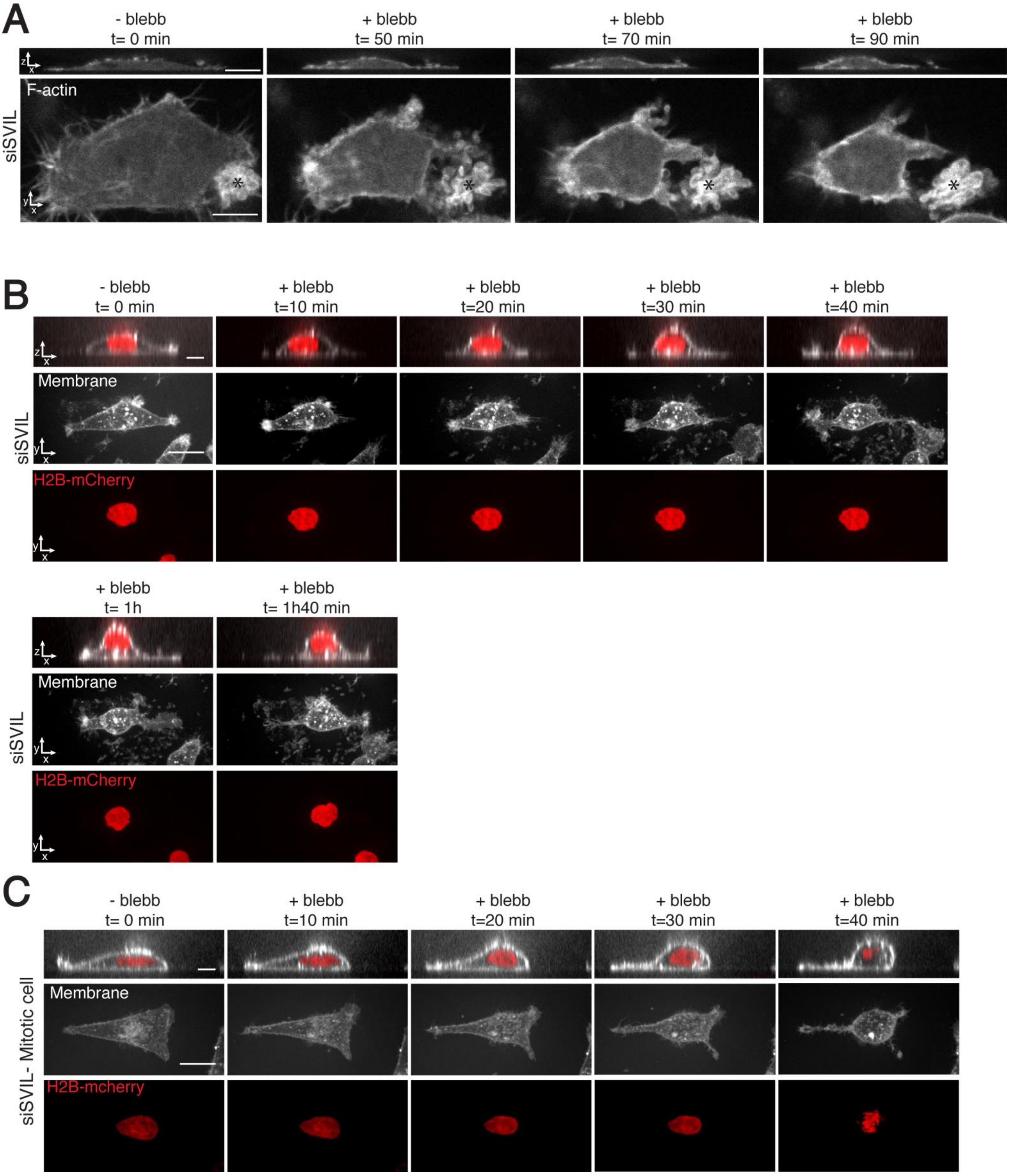
A
- blebb
t= 0 min
+ blebb
t= 50 min
+ blebb
t= 70 min
+ blebb
t= 90 min
siSVIL
F-actin
B
- blebb
t= 0 min
+ blebb
t=10 min
+ blebb
t=20 min
+ blebb
t=30 min
+ blebb
t=40 min
Membrane
siSVIL
H2B-mCherry
+ blebb
t= 1h
+ blebb
t= 1h40 min
Membrane
siSVIL
H2B-mCherry
C
- blebb
t= 0 min
+ blebb
t=10 min
+ blebb
t=20 min
+ blebb
t=30 min
+ blebb
t=40 min
Membrane
siSVIL- Mitotic cell
H2B-mcherry

**Figure S4: Control experiments for the effect of supervillin depletion and myosin activity inhibition on cell shape, related to Figure 3.**

**(A)** Representative time lapse images of supervillin-depleted cells expressing GFP-Lifeact before and after addition of blebbistatin. Note the increase in cortical F-actin over time. The asterisks indicate a dead cell. Scale bars XY, 10 µm, XZ, 10 µm. **(B)** Representative time lapse images of H2B-mCherry-expressing HeLa cells treated with siRNA against supervillin (siSVIL), and with the myosin II inhibitor blebbistatin at t=0. Far red cell mask was used to label the membrane. Note that cell rounding occurs in interphase, as evidenced by DNA appearance. Scale bars XY, 20 µm, XZ, 10 µm. **(C)** Representative time lapse images of H2B-mCherry-expressing HeLa cells treated with siRNA against supervillin (siSVIL) and entering mitosis, as evidenced by DNA condensation. Far red cell mask was used to label the membrane. Scale bars XY, 20 µm, XZ, 10 µm.
